# Performance of Reasoner’s 2 Agar (R2A) medium in enumeration of culturable microorganisms in water intended for human consumption

**DOI:** 10.64898/2026.02.09.702495

**Authors:** Kristiina Valkama, Anna Pursiainen, Olivier Molinier, Sunna Nikodemus, Eric Pierlot, Tarja Pitkänen

**Author notes:** Corresponding author: Tarja Pitkänen.

## Abstract

Enumeration of the heterotrophic plate counts of culturable microorganisms (HPC) is used for the assessment of water quality before, during and after the drinking water treatment processes and monitoring of integrity of the drinking water distribution systems. Any substantial change in HPC is a warning of potential contamination, treatment failure, or intrusion. This study produced data for international standardization of Reasoner’s 2 Agar (R2A) method. Interlaboratory studies were applied to find out which inoculation technique, spread plate or pour plate, is more applicable, and what is the acceptable transport and storage time between the sampling and initiation of the analysis. Based on the produced interlaboratory data, spread plate inoculation followed by incubation for 7 days at 22 °C was selected as the standard method, and it is recommended to analyse samples as soon as possible, but the samples may be kept at (5 ± 3) °C for up to 24 hours after the sampling prior to examination. Performance characteristics of the method were determined in a single laboratory according to ISO 13843:2017 by using process water from waterworks, bottled water, chlorinated and non-chlorinated tap water, and well water intended for human consumption. For quality assurance of the R2A medium, the use of control strains of *Bacillus subtilis subsp. spizizenii*, *Pseudomonas fluorescens* and an additional strain of *Sphingomonas paucimobilis* was verified during the work. The R2A method is especially suitable for determination of micro-organisms forming colonies after a prolonged incubation time, enabling HPC enumeration from biostable waters with low nutrient levels and low temperature. Such waters usually produce zero counts when nutrient-rich formulations of culture media with short incubation times are employed. Further, the nutrient limited conditions of R2A minimize the colony size and overgrowth in all kinds of water providing improvement to the HPC determinations.

## 1. Introduction

Heterotrophic plate count (HPC) is the colony count of all micro-organisms (bacteria, yeasts and moulds) forming visible colonies by a particular method, specified with inoculation technique, composition of the culture medium, and time and temperature of incubation (Allen et al., 2004; ISO 6222, 1999). The enumeration of the overall colony count of culturable microorganisms provides useful information for the assessment and surveillance of the raw water quality, efficiency of water treatment processes and integrity of the drinking water distribution systems. HPC contains a variety of microorganisms derived from various sources, such as raw water resource, different treatment steps within drinking water production and drinking water distribution networks. The levels of HPC reflect the health of the distribution network with connection to the availability of nutrients and biostability, rather than having direct relation to human health (Allen et al., 2004; Carabin et al., 2024). However, among the heterotrophic microorganisms, also opportunistic pathogens occur and high HPC in tap water may cause a health risk especially for immuno-compromised individuals (Pavlov et al., 2004).

A recent comprehensive review gave a reminder to water practitioners that while HPC is widely assessed in drinking water distribution systems, methodological standards for HPC determinations are not clearly defined (Carabin et al., 2024). Deviation in the methodology is evident in the field, for example in the recent literature, Faraji et al. (Faraji et al., 2025) employed pour plating technique for Reasoner’s 2 Agar medium (R2A) for 72 hours at 35°C, Rygala et al. (Rygala et al., 2020) utilized membrane filtration technique followed by incubation on R2A for 3 days at 22 °C and for 2 days at 37 °C, and Sigudu et al. (2024) used spread plating technique on a nutrient agar plates that were incubated for 48 h at 35 °C, for HPC analysis. It is well known that HPCs differ significantly when compared between different culture media and incubation conditions (Pitkänen et al., 2011; Reasoner et al., 1989). While the available methodological options provide great flexibility in the application of the HPC analysis to drinking water (Reasoner, 2004), the deviations in the HPC enumeration methods make it impossible to compare the HPC levels between the studies.

There is a need for method standardization, since the main value of colony counts lies in the detection of changes from those expected, based on representative monitoring. The change in colony counts can be caused by various reasons like biofilm formation, increase in water temperature, presence of microbially available nutrients or decrease in chlorine concentration (reviewed by Carabin et al., 2024). Any substantial increase in count can also be either a warning of potential treatment failure or contamination. This indication value of HPC is considered in guidelines for drinking water safety, such as the European Union Drinking Water Directive (EU DWD 2020/2184) with specification that there should be no abnormal changes in colony counts at 22 °C.

Both high-nutrient and low-nutrient media are used for HPC determinations. Nutrient rich tryptone yeast extract agar medium has been used for long time in water microbiology (ISO 6222, 1999). ISO 6222 is the reference method of EU DWD (2020/2184) and it specifies the detection of microorganisms in nutrient rich conditions at 22 °C in 3 days or at 36 °C in 2 days. However, the use of Reasoner’s 2 Agar (R2A) medium (Reasoner & Geldreich, 1985) that has a low-nutrient composition and long incubation time (7 days at 22 °C) provides an improvement for the enumeration of autochthonous microorganisms found in aquatic systems including drinking water distribution systems (Allen et al., 2004). The R2A method is considered especially suitable for providing more information for monitoring water for human consumption in waters with low colony counts, i.e., waters low in nutrients and is distributed in temperatures below 20 °C (Pitkänen et al., 2011). The R2A method usually gives higher colony counts from water samples than typically used nutrient-rich formulations of culture media (Allen et al., 2004).

The aim of this work was to validate the R2A method for enumeration of heterotrophic plate counts of culturable microorganisms (HPC) from water by spread plate technique to serve as a new International Standard ISO 13647 (Water quality — Enumeration of culturable microorganisms — Colony count by spread plate inoculation on R2A medium). There are three inoculation techniques in use in water microbiology (ISO 8199, 2018; Lipps et al., 2004): spread plate technique (typically used with 0,1 ml sample volumes), pour plate technique (typically used with 1 ml sample volumes) and membrane filtration (sample volume varies but is typically 100 ml). In determinations of heterotrophic plate counts, all these three inoculation methods have been applied with different culture media including R2A. To reach consensus regarding the most useful inoculation technique, spread plate and pour plate techniques on R2A were compared. Further, plate counts on R2A were compared to counts on tryptone yeast extract agar (ISO 6222, 1999). Performance characteristics of the R2A method and robustness of incubation time and temperature were determined and the acceptable storage time between sampling and initiation of the R2A analysis was verified.

## 2. Materials and methods

### 2.1. Water samples

The water samples used for the method validation purposes in this study represented natural conditions in the drinking water systems, i.e., the samples were studied as they are without any spikes of microbes or other modifications. In the experiment comparing the spread plate and pour plate techniques, surface water, spring water, well water, ground water, process water from the drinking water production, water from drinking water distribution system, water storage reservoir and tap water were investigated in 11 laboratories from six countries (**Table 1a**).

**Table 1.** Origin of samples (country), number of laboratories, water types and number of water samples (N) in **a)** the experiment comparing the spread plate and pour plate techniques, **b)** the single laboratory study determining the R2A method performance characteristics, and **c)** the study determining acceptable storage time between sampling and initiation of the R2A analysis.

| Origin | Laboratories | Water type | N |
| --- | --- | --- | --- |
| *A. Comparison between the inoculation techniques* |  |  |  |
| Austria | 1 | Surface water | 1 |
|  |  | Spring water | 2 |
|  |  | Ground water | 2 |
|  |  | Drinking water: distribution system | 8 |
|  |  | Drinking water: water storage reservoir | 1 |
|  |  | Drinking water: tap water | 2 |
| Egypt | 1 | Surface water | 3 |
| Finland | 3 | Drinking water: well water | 5 |
|  |  | Drinking water: distribution system | 2 |
|  |  | Drinking water: tap water | 33 |
| Germany | 3 | Surface water | 1 |
|  |  | Drinking water: well water | 15 |
|  |  | Process water from the drinking water production | 23 |
|  |  | Drinking water: distribution system | 15 |
|  |  | Drinking water: tap water | 17 |
| Netherlands | 1 | Ground water | 1 |
|  |  | Drinking water: distribution system | 19 |
| Sweden | 2 | Process water from the drinking water production | 10 |
|  |  | Drinking water: tap water | 30 |
| **Total** | **11** |  | **190** |
| *B. Single laboratory performance characteristics study* |  |  |  |
| Finland | NA | Spring water | 1 |
|  |  | Drinking water: well water | 17 |
|  |  | Drinking water: water storage reservoir | 19 |
|  |  | Drinking water: bottled water | 1 |
|  |  | Drinking water: tap water | 57 |
| Norway | NA | Drinking water: bottled water | 2 |
| **Total** |  |  | **97** |
| *C. Study of acceptable storage time prior analysis* |  |  |  |
| Austria | 1 | Ground water | 12 |
| England | 1 | Ground water | 3 |
|  |  | Process water from the drinking water production | 5 |
|  |  | Drinking water: distribution system | 4 |
|  |  | Drinking water: tap water | 1 |
| Finland | 1 | Ground water | 1 |
|  |  | Process water from the drinking water production | 3 |
|  |  | Drinking water: tap water | 4 |
| France | 2 | Spring water | 1 |
|  |  | Drinking water: tap water | 7 |
|  |  | Water used in hospital | 30 |
| Germany | 2 | Drinking water: tap water | 6 |
|  |  | Drinking water: well water | 2 |
|  |  | Drinking water: distribution system | 10 |
| Netherlands | 2 | Drinking water | 55 |
| Portugal | 1 | Process water from the drinking water production | 15 |
|  |  | Drinking water: tap water | 4 |
| Republic of Korea | 2 | Raw water | 4 |
|  |  | Process water from the drinking water production | 6 |
|  |  | Drinking water: bottled water | 1 |
|  |  | Drinking water: tap water | 5 |
| Slovakia | 1 | Raw water | 2 |
|  |  | Process water from the drinking water production | 2 |
|  |  | Drinking water: distribution system | 2 |
|  |  | Drinking water: tap water | 4 |
| **Total** | **13** |  | **198** |

NA, not applicable.

To determine the performance characteristics of the R2A method and robustness of incubation time and temperature, spring water, well water, water storage reservoir and tap water were collected from the locations nearby the city of Kuopio, Finland. Bottled water from both Finland and abroad was also used to determine the performance characteristics of the R2A method (**Table 1b**). All water samples investigated by the water microbiology laboratory of the Finnish Institute for Health and Welfare.

In the study determining acceptable storage time between sampling and initiation of the R2A analysis, raw water, spring water, well water, ground water, process water from the drinking water production, water from drinking water distribution system, bottled water, tap water and water used in hospital were investigated in 13 laboratories from nine countries (**Table 1c**).

### 2.2. Culture media

Preparation of all culture media was carried out according to the manufacturer’s instructions. The laboratories prepared the R2A agar plates in-house or used ready-made agar plates in the experiments.

The R2A culture media used in the comparison of the spread plate and pour plate techniques were manufactured by Oxoid (United Kingdom), Merck (Germany), Roth (Germany), Neogen (United States), BD (United States) and Difco (United States).

For determination of the performance characteristics of the R2A method, R2A was produced from Reasoner’s 2 Agar (R2A), BD 218263 (BD, United States), tryptone yeast extract agar from water plate count agar, CM1012 (Oxoid, United Kingdom) and tryptone soya agar (TSA) from tryptone soya agar, CM0131 (Oxoid, United Kingdom).

In a determination of an acceptable storage time between sampling and initiation of the R2A analysis, laboratories used R2A obtained from BD (United States), Oxoid (United Kingdom), SGL (United Kingdom), Difco (United States), Thermofisher (United States), Merck (Germany), Roth (Germany) and Merck Millipore (Germany).

### 2.3. Determining the inoculation technique

The comparison of the spread plate and pour plate techniques was conducted as an interlaboratory study which involved 11 laboratories from six countries. The aim was to find out if there was a difference in the HPC or dispersion of HPC when samples were analysed in different laboratories using spread plate technique on Reasoner’s 2 Agar (R2A) and pour plate technique on tryptone yeast extract agar (ISO 6222, 1999). To investigate this, three parallel plates were cultivated from each water sample using spread plate and pour plate techniques. In both techniques, 0.1 ml of diluted or undiluted sample was analysed per plate and plates were incubated at 22 ± 2 °C 164 ± 4 hours. The diluents were prepared and the inoculations performed by following the instructions of ISO 8199 (2018). After incubation, the total number of colonies present in each plate were counted immediately by using magnifying equipment. If dilutions were used, result as colony forming unit (CFU)/ml were calculated using weighted mean according to ISO 8199 (2018).

### 2.4. A single laboratory validation study

The effect of diluents on HPC as well as the effect of magnification and location of light source on colony counting was investigated in a single laboratory study at the Finnish Institute for Health and Welfare, Kuopio, Finland. Further, the performance characteristics of R2A method were determined and all calculations were conducted in accordance with ISO 13843 (2017). Determined performance characteristics were sensitivity, proportion of false positive results, upper limit, repeatability, reproducibility, relative recovery, uncertainty of counting, and the robustness of the incubation time and temperature. The single laboratory validation study was conducted using the spread plate inoculation method and the colonies were counted from plates according to the principles of ISO 8199 (2018). Unless weighted mean was used or stated otherwise, plates with colony count of 4-200 were included in the study.

#### 2.4.1. Effect of diluents on HPC

Sterile deionized water and diluents described in ISO 8199 (2018) including physiological saline solution (Merck, Denmark), phosphate buffer (Merck, Germany), peptone water (Oxoid, United Kingdom) and peptone saline solution (Oxoid, United Kingdom and Merck Denmark) were used to determine how diluents used in preparation of the samples and during the inoculation affect the performance of the R2A method. All tested diluents were manufactured at the water microbiology laboratory of the Finnish Institute for Health and Welfare in accordance with ISO 8199 (2018).

The test of the diluents was conducted according to ISO 11133 (2014). Dilutions were prepared by pipetting 1 ml of sample into a tube, which contained 9 ml of the selected diluent. A volume of 0.1 ml of each dilution was spread plated on R2A medium using each diluent to be compared. Each dilution was cultivated on two parallel plates directly after dilution. After the first cultivation, the tubes were left for 45 minutes on the laboratory bench before re-cultivation on two parallel plates to determine the effect of diluent on the plate counts. The dilutions were tested with nine water samples including well water and tap water.

#### 2.4.2. Magnification and location of light source in colony counting

To study how magnification used in colony counting affects the performance of the R2A method, colonies on the R2A plates were counted by using magnifications of 0x (naked eye), 1.85x Luxo Wave magnifying lamp (Glamox Luxo, Norway), 2.25x Solar Comp LED lamp and 3x Schuett Count colony counter (Schuett-Biotec, Germany). As a reference method, colonies were counted also with an Olympus SZH-ILLB stereo microscope (Olympus Corporation, Japan). Same plates were counted with all magnifications moving from the lowest to the highest magnification by five operators. Eleven water samples originating from drinking water distribution systems, taps and wells were used and this produced results from a total of 56 plates.

The impact of the location of the light source on colony counting was tested by counting the colonies on the R2A plates using a light located above the plate and below the plate by five operators. A dark background and magnifications 2.25x and 3x were used for counting. A total of 24 plates containing colonies from drinking water distribution systems and tap water were used.

#### 2.4.3. Sensitivity and proportion of false positive results

Sensitivity is proportion of the total number of colonies correctly identified during the presumptive counting. In HPC determinations, false positive results refer to objects that are initially counted but are subsequently not confirmed of being colonies of microorganisms. In the determination of the sensitivity and proportion of false positive results, colonies on R2A plates were counted by one operator using 2.25x magnification and a light source located above the plate and then verified with a stereo microscope. Thirteen water samples were used including well water, spring water, water storage reservoir and tap water, and a total of 24 plates that each contained 20-200 colonies were examined. True positive, false positive and false negative colonies on the R2A plates were recorded.

#### 2.4.4. Upper limit

Upper limit is the highest applicable limit of the method’s working range, in case of R2A the maximum number of countable colonies in one solid agar medium plate using a 90 mm Petri dish. To determine the upper limit of the method, a total of 10 water samples were used including well water, spring water, water storage reservoir and tap water. The upper limit was determined with the samples which estimated plate count exceeded the expected upper limit of 200 colonies. Two-fold dilution series with six successive steps were prepared, and three parallel plates of each dilution level were cultivated. Proportionality of the obtained colony counts was evaluated as the log-likelihood ratio statistic (G^2^ test) according to ISO 13843 (2017). If the linearity was not acceptable, the dilution levels exhibiting non-linear counts were removed and the value of the index was compared again with the χ2 distribution with n-1 degrees of freedom which depends on the number of dilution levels (ISO 13843, 2017). This allowed to determine the highest reliable number of colonies per plate.

#### 2.4.5. Repeatability and reproducibility

Repeatability is precision of the method obtained during repeatability conditions such as same operator and same operating conditions in a short period of time. In the determination of the repeatability, one operator cultivated 10 parallel R2A plates of the same sample in as short time as possible and in identical conditions. Same operator counted the colonies on the R2A plates and after that overdispersion was calculated. The test of repeatability was performed by two different operators. One operator cultivated and counted seven water samples, and the other four water samples. Used water samples included well water and tap water.

Reproducibility of cultivation is precision of the method during reproducibility conditions such as different operator, different operating conditions and replicate measurements from the same sample. Three to four operators cultivated samples in different laboratory conditions such as different R2A plate batches, cultivation equipment and cultivation bench to determining the intralaboratory reproducibility of cultivation. A total of nine water samples were used including well water, spring water and tap water. Colony counting from all the R2A plates was performed by a single operator, so only the repeatability of the cultivation conditions was examined in analysis of the relative operational variance of colony counts on R2A plates.

#### 2.4.6. Relative recovery

Relative recovery describes how effectively method detects the target organisms from the sample compared to another method. The relative recovery of the R2A method was determined at 22 ± 2 °C by comparing the colony counts obtained by the spread plate technique on R2A agar medium after 7 days to the colony counts after 3 days using pour plate technique on nutrient rich tryptone yeast extract agar (ISO 6222, 1999). A total of 15 water samples were analysed including well water, spring water and tap water. The water samples with plates up to 200 colonies were included in the comparison. The relative recovery was determined in accordance with ISO 17994 (2014).

#### 2.4.7. Uncertainty of counting

Uncertainty of counting is relative standard deviation of repeated counting of the colonies under specified conditions such as same operator or between different operators in a same laboratory. Personal uncertainty of counting for each operator and intralaboratory uncertainty of counting were calculated.

The same operator counted the colonies on the R2A plates twice to determine personal uncertainty of counting. The number of plates counted ranged between five and 30 in the determination of the personal uncertainty of counting. In the determination of the intralaboratory uncertainty of counting, a total of 30 plates with colony counts ranging from 20 to 200 and originating from 17 different water samples were counted by eight operators. Used water samples included well water, spring water, water storage reservoir and tap water.

#### 2.4.8. Robustness of incubation time and temperature

Robustness of incubation time and temperature were determined to investigate how incubation time (164 ± 4 hours) and incubation temperature (22 ± 2 °C), as described in ISO 13647 (2026), affect to the colony counts on R2A. For robustness of incubation time, samples were diluted and cultivated as three parallel plates and plates were incubated at 22 ± 2 °C. Plates were counted after incubation of 160 hours, 164 hours and 168 hours by two operators. To determine robustness of incubation time, a total of 11 samples were used including well water, bottled water and tap water.

For robustness of incubation temperature, samples were diluted and cultivated as three parallel plates and plates were incubated at 20.5 °C and 23.5 °C for 164 ± 4 hours, and a total of seven samples were used including well water, bottled water and tap water. Relative recovery between different incubation times and incubation temperatures for separate plates were calculated in accordance with ISO 17994 (2014).

### 2.5. Determining acceptable storage time between sampling and initiation of the analysis

Acceptable time between sampling and initiation of R2A analysis were determined by triplicate analysing of water samples before and after storage in 5±3 °C in 13 laboratories from nine countries by using a total of 198 samples (**Table 1c**). Each laboratory analysed a minimum of eight routine samples. Spread plate inoculation of 0,1 ml was applied successfully for 176 samples after storage up to 12 h and at least 24 h following sampling, corresponding to 168 statistically processable pairs of data (removing the 8 samples with results including a zero).

### 2.6. Control strains for quality assurance of the R2A medium

Performance testing for the quality assurance of the R2A medium were conducted with four control strains *Pseudomonas fluorescens* WDCM 00115, *Bacillus subtilis subsp. spizizenii* WDCM 00003, *Sphingomonas paucimobilis* WDCM 00232 and *Sphingomonas echinoides* DSM 1805. All four control strains were first tested in one laboratory. After that, *Pseudomonas fluorescens* WDCM 00115, *Bacillus subtilis subsp. spizizenii* WDCM 00003 and *Sphingomonas paucimobilis* WDCM 00232 were tested in another laboratory in a different country. Each control strain was cultivated on two or three parallel plates and plates were incubated at 22 ± 2 °C three days for *Pseudomonas fluorescens* and *Bacillus subtilis subsp. spizizenii* and seven days for *Sphingomonas paucimobilis* and *Sphingomonas echinoides*. Poisson index of dispersion between counts on parallel R2A plates was calculated according to ISO 13843 and if dispersion was acceptable, productivity ratio (Pr) was calculated for each control strain following the principles of ISO 11133 (2014) by using an earlier batch of R2A medium as the reference medium, i.e. the same medium was used as the reference medium as the medium under the test. Another R2A batch represented a non-selective reference medium in the calculations since in case of R2A agar there is no other medium which enables the growth of the same heterotrophic biota as R2A agar medium.

### 2.7. Statistical analyses

A p-value of less than 0,05 was used to indicate that there is a statistically significant difference between the distributions of the data tested. For determining the performance characteristics of the R2A method, Poisson index of dispersion and determination of relative operational variance were utilized following the principles of ISO 13843 (2017).

A non-parametric Wilcoxon test for paired series based on rank was used to determine the statistical difference in the comparison of the spread plate and pour plate techniques and in the determination of the robustness of incubation time and temperature. Statistical differences between diluents, magnifications and location of the light were determined with a non-parametric Kruskal-Wallis test.

Two complementary statistical approaches were used to evaluate the paired data between 12 and 24 hours in statistical analyses in the determination of the acceptable time between sampling and initiation of R2A analysis. Wilcoxon test, with no pre-requisites on the distribution of results, and the calculation of relative recovery of microorganisms by two storage times and to assess the order of magnitude of the deviation in accordance with ISO 17994 (2014), considering for the calculations that 12-h and 24-h storages were two different methods, were used. Additional statistical processing was performed from each laboratory storage times data using adapted ISO 17994 approach. The adjustment involved a coverage factor based on the Student distribution adapted to the small number of data.

## 3. Results

### 3.1. Comparison of the spread plate and pour plate techniques

A total of 190 samples were analysed in 11 laboratories to compare spread plating and pour plating techniques for HPC determinations on R2A medium. The spread plate technique resulted in a higher HPC from 159 samples (84%) and pour plate technique from 30 samples (16%) (**Figure 1**). Spread plate and pour plate techniques gave the same result for one sample. On average, the HPC was significantly higher with the spread plate technique than pour plate technique (**Supplemental Table S1**; Wilcoxon: p < 0.001). However, also the dispersion between three replicate plates was higher with a spread plate technique than pour plate technique. The square root of relative operational variance, i.e. repeatability, was for spread plate technique 17% and for pour plate technique 5%.

**Figure 1.**
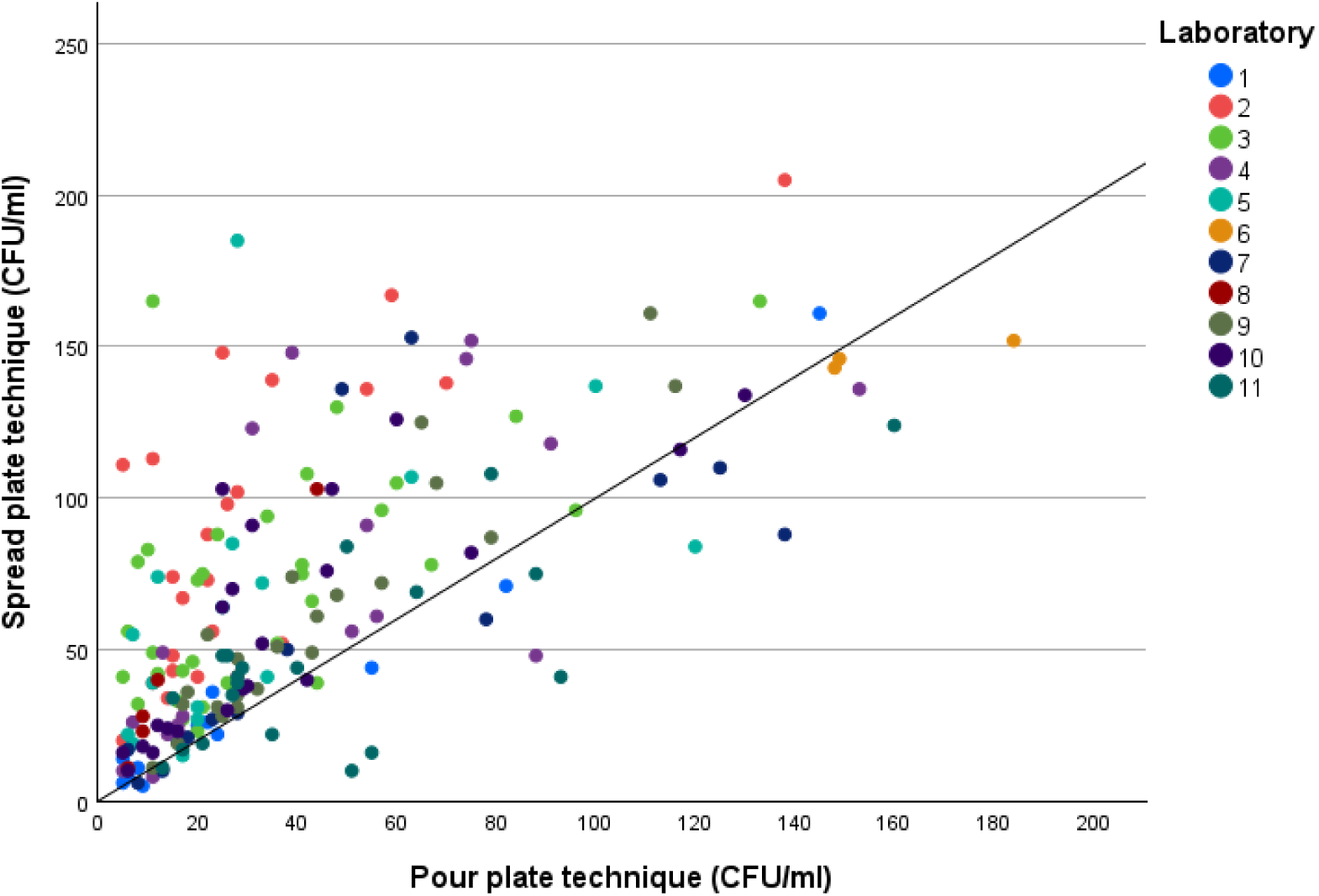
Results of the comparison between spread plate and pour plate techniques for inoculation of water samples on R2A agar medium in 11 participating laboratories. Each point represents the arithmetic mean of three parallel plates of the sample and the straight line represents the case when colony count (CFU/ml) by using spread plate and pour plate techniques would have been equal (x=y).

### 3.2. Effect of diluents on HPC

In the comparison of diluents, nine samples were analysed. The average percentage difference between the samples’ plate count after incubation of the 10-fold dilution series for 0 minutes and 45 minutes prior spread plating on R2A medium exceeded the 30% limit allowed by ISO 11133 (2014) when physiological saline and peptone saline solution were used as the diluents (**Table 2**). Based on the arithmetic mean values, highest plate counts were detected by using sterile water, whereas the lowest plate counts detected by using peptone saline solution although the difference between the diluents was not statistically significant (Kruskal-Wallis: p=0.718). Sterile water, physiological saline, phosphate buffer and peptone saline solution decreased the colony count in 45 minutes, whereas dilution with peptone water increased the colony count (**Supplemental Table S2**). However, the difference between the inoculation time points of 0 min and 45 min was not statistically significant with any diluent (Mann-Whitney U test: sterile water p=0.800; physiological saline p=0.268; phosphate buffer p=0.624; peptone water p=0.501; peptone saline solution p=0.226).

**Table 2.** Changes in the plate counts of the nine studied samples in dilution solutions after 0 min and 45 min on R2A medium. The percentage change has been calculated from the arithmetic mean of two parallel plates.

| Sample | Sterile water (%) | Physiological saline (%) | Phosphate buffer (%) | Peptone water (%) | Peptone saline solution (%) |
| --- | --- | --- | --- | --- | --- |
| 1 | 42.0 | -27.3 | 22.0 | 10.0 | -43.2 |
| 2 | 5.9 | -20.8 | -21.2 | -16.7 | -25.9 |
| 3 | -9.6 | -27.2 | -13.7 | 0.6 | 8.3 |
| 4 | -52.4 | -12.5 | 0.0 | 29.4 | -75.0 |
| 5 | -110.0 | -35.7 | -82.4 | -23.5 | 21.4 |
| 6 | -13.5 | -16.7 | -50.0 | 47.5 | -104.3 |
| 7 | -25.1 | -44.5 | -21.1 | 5.3 | -31.6 |
| 8 | -3.6 | -78.9 | 20.6 | 20.6 | -5.3 |
| 9 | -0.7 | -7.4 | -3.7 | 15.9 | -71.9 |
| **Arithmetic mean** | **-18.6** | **-30.1** | **-16.6** | **9.9** | **-36.4** |

### 3.3. Magnification and location of light source in colony counting

In the first comparison, 16 plates were counted using magnifications of 0x (naked eye), 1.85x, 2.25x, 3x and with a stereo microscope, which was the reference method. On the average, the highest colony counts were detected using magnification of 2.25x and the lowest colony counts detected with naked eye (**Table 3a**). Colony counts detected using magnification of 2.25x were also closest to the counts obtained by using stereo microscope. The difference between magnifications was not statistically significant (Kruskal-Wallis p=0.596). In the second comparison, magnifications of 2.25x, 3x and a stereo microscope were compared by counting 40 plates. Higher colony counts were detected using magnification of 2.25x than magnification of 3x. Also, the colony counts detected using magnification of 2.25x were closer to the reference method than magnification of 3x (**Table 3b**). However, the difference between magnifications of 2.25x and 3x was not statistically significant (Mann-Whitney: p=1.96). In addition, there was no statistically significant difference between the magnifications of 2.25x and 3x and the stereo microscope (Kruskal-Wallis p=0.381).

**Table 3.** Sum of colonies and descriptive statistics of HPC with a stereo microscope and magnifications of a) 0x (naked eye), 1.85x, 2.25x, and 3x and b) 2.25x and 3x. ND, not done.

| Magnification | 0x | 1.85x | 2.25x | 3x | Stereo microscope |
| --- | --- | --- | --- | --- | --- |
| *a) 1<sup>st</sup> comparison (N=16 plates)* |  |  |  |  |  |
| Sum of colonies | 2350 | 2558 | 2755 | 2634 | 2673 |
| Arithmetic mean | 53 | 58 | 63 | 60 | 62 |
| Median | 50 | 56 | 56 | 55 | 61 |
| Standard deviation | 31 | 34 | 35 | 33 | 33 |
| *b) 2<sup>nd</sup> comparison (N=40 plates)* |  |  |  |  |  |
| Sum of colonies | ND | ND | 6607 | 5856 | 6485 |
| Arithmetic mean | ND | ND | 51 | 46 | 51 |
| Median | ND | ND | 37 | 33 | 36 |
| Standard deviation | ND | ND | 39 | 36 | 39 |

No statistically significant differences (Kruskal Wallis p=0.465) were observed in the comparison of the location of the light during colony counting when 24 plates were counted with magnification of 2.25x and 3x light source below and above the plate and with stereo microscope (**Supplemental Table S3**).

### 3.4. Performance characteristics of the R2A method

#### 3.4.1. Sensitivity and proportion of false positive results

To determine the sensitivity and the proportion of false positive results, the total number of true positives, false positives and false negatives on 24 plates were used (**Table 4**).

**Table 4.** Total number of true positives, false negatives and false positives used to determine sensitivity and the proportion of false positives (N=24 plates).

|  |  | Initially counted colonies |  |
| --- | --- | --- | --- |
|  |  | + | - |
| Confirmed colonies | + | True positives: 1520 | False negatives: 35 |
|  | - | False positives: 45 | True negatives: - |

Sensitivity was determined based on the sum of true positive and false negative colonies, and the result for the sensitivity was 98%. The proportion of false positives was determined based on the sum of true positives and false positives, the result for the false positives was 3%.

#### 3.4.2. Upper limit

In determination of the upper limit for HPC on R2A, in total of 10 samples were analysed. Except for one sample, for all samples proportionality (G^2^ test value) was not at an acceptable level when the average HPC exceeded 200 on the least diluted level (**Table 5**). In those occasions, linearity was reached after removing the dilution levels resulting in exceedances.

**Table 5.**
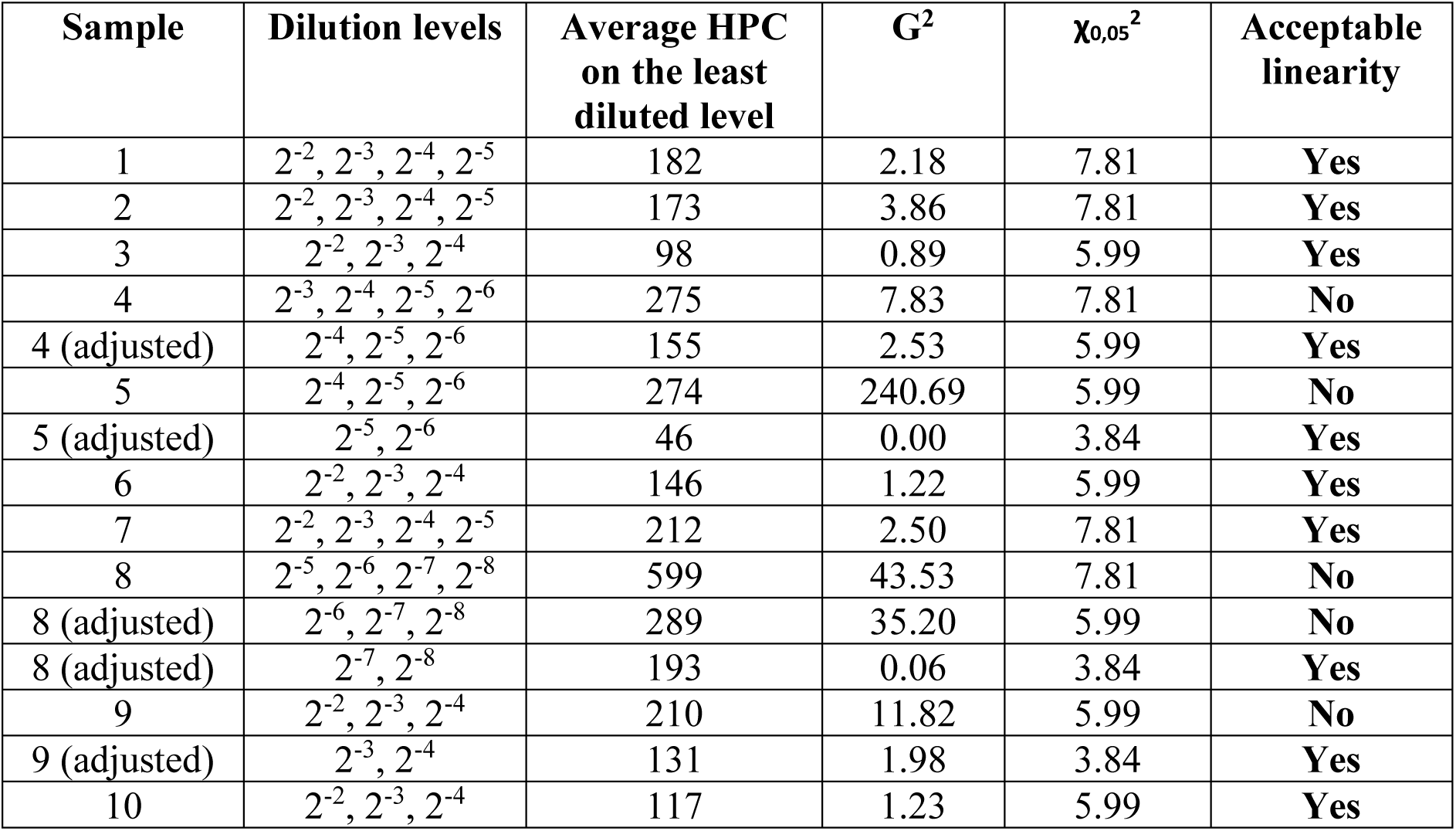
Dilution levels of the 10 samples studied to determine the upper limit for plate count, arithmetic mean of plate counts on parallel plates (n=3) on the least diluted level, G^2^ value between the levels of dilution and comparison of the linearity with Poisson distribution index (χ0.05^2^). Dilution levels included in calculations were adjusted where needed to reach acceptable linearity.

#### 3.4.3. Repeatability and reproducibility

Repeatability was determined from seven samples, which were cultivated as 10 parallel plates. Only in one sample, the deviation between the replicates was too large compared to Poisson distribution index (ISO 13843:2017) (International Organization for Standardization, 2017) (**Table 6a**). The mean relative operative variance of the samples was 0.004 and square root of relative operational variance, i.e. repeatability, was 6%.

**Table 6.** The lowest and highest plate count, median, arithmetic mean and relative operational variance (uo^2^) of a) 10 replicates used for the determination of repeatability, where dispersion between parallel plates (χ ^2^) was compared with the value of Poisson distribution index 16.919 to determine if the dispersion was acceptable, and b) duplicates plated by different operators for determination of reproducibility, NA, not applicable.

| Sample | Operators | Min | Max | Median | Arithmetic mean | $u_o^2$ | $\chi_{r-1}^2$ | Dispersion between parallel plates |
| --- | --- | --- | --- | --- | --- | --- | --- | --- |
| *a) Repeatability test (N=7)* |  |  |  |  |  |  |  |  |
| 1 | 1 | 51 | 71 | 59 | 60 | -0.005 | 6.072 | Accepted |
| 2 | 1 | 56 | 79 | 74 | 72 | -0.004 | 6.262 | Accepted |
| 3 | 1 | 13 | 29 | 17 | 19 | 0.051 | 17.789 | Not accepted |
| 4 | 1 | 66 | 99 | 91 | 87 | 0.003 | 11.211 | Accepted |
| 5 | 1 | 9 | 16 | 13 | 12 | -0.032 | 5.529 | Accepted |
| 6 | 1 | 83 | 111 | 92 | 94 | 0.000 | 8.733 | Accepted |
| 7 | 1 | 8 | 21 | 16 | 16 | -0.001 | 8.872 | Accepted |
| *b) Reproducibility test (N=9)* |  |  |  |  |  |  |  |  |
| 1 | 4 | 67 | 98 | 81 | 82 | 0,003 | NA | NA |
| 2 | 4 | 16 | 33 | 23 | 24 | 0,023 | NA | NA |
| 3 | 4 | 74 | 94 | 78 | 81 | -0.003 | NA | NA |
| 4 | 3 | 20 | 35 | 28 | 28 | -0.004 | NA | NA |
| 5 | 3 | 101 | 186 | 158 | 154 | 0.029 | NA | NA |
| 6 | 3 | 2 | 19 | 11 | 10 | 0.190 | NA | NA |
| 7 | 3 | 67 | 88 | 76 | 76 | -0.002 | NA | NA |
| 8 | 3 | 120 | 159 | 130 | 137 | 0.009 | NA | NA |
| 9 | 3 | 44 | 62 | 50 | 53 | 0.002 | NA | NA |

Reproducibility was determined from nine samples, which were cultivated as two parallel plates by three to four operators (**Table 6b**). The mean relative operative variance of the samples was 0.027 and the square root of relative operational variance, i.e. reproducibility, was 17%.

#### 3.4.4. Relative recovery

To determine relative recovery, 15 samples were analysed, and samples were cultivated as duplicate plates with both spread plate technique on R2A and pour plate technique on tryptone yeast extract agar. The weighted mean of the duplicate results per sample was calculated from parallel plates (**Figure 2**). The relative difference according to ISO 17994 (2014) between spread plate technique on R2A and pour plate technique on nutrient rich tryptone yeast extract agar was 300%.

**Figure 2.**
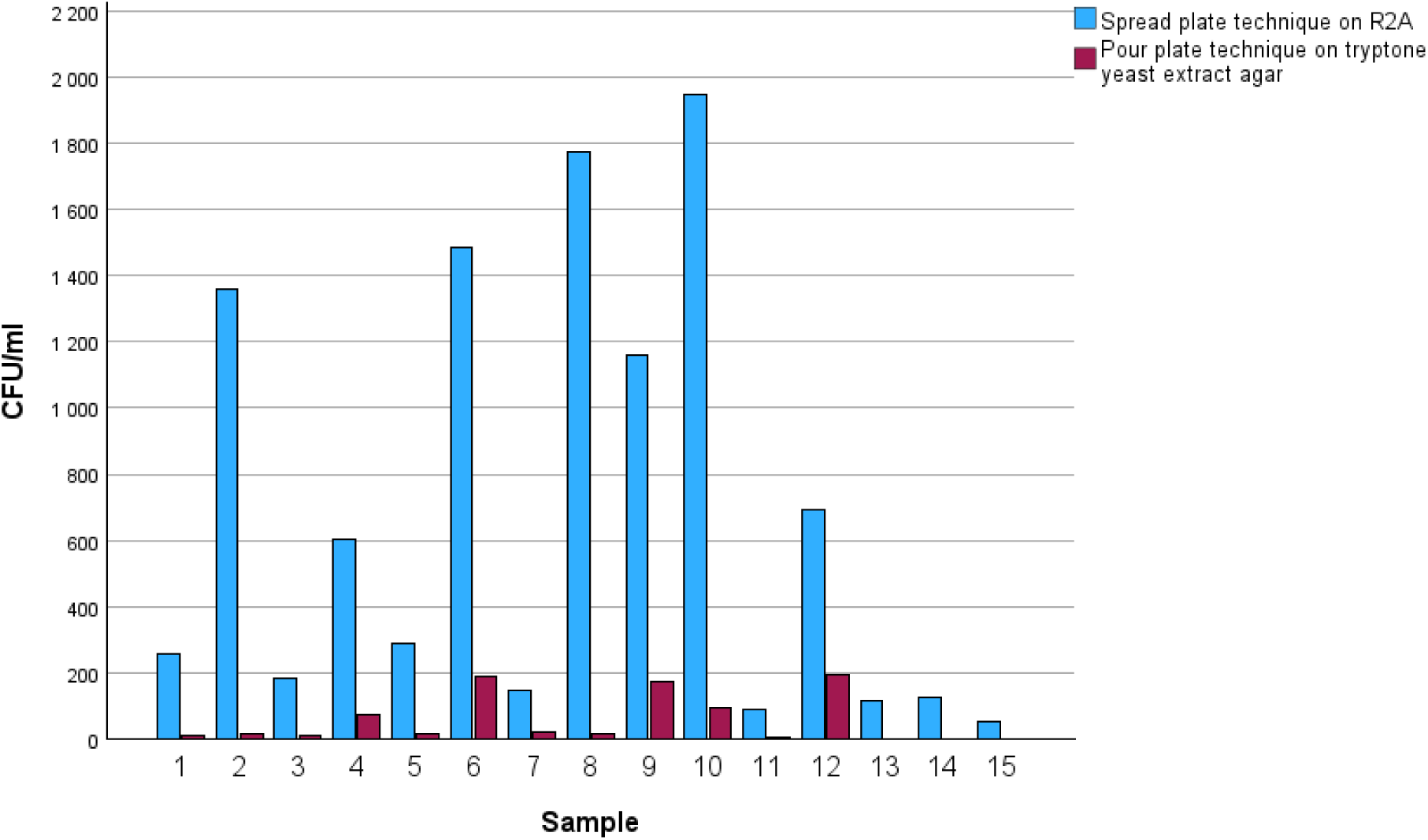
Number of colonies (CFU/ml) in water intended for human consumption analysed by spread plate technique on R2A and pour plate technique on nutrient rich tryptone yeast extract agar.

#### 3.4.5. Uncertainty of counting

In determination of the personal and intralaboratory uncertainty of counting, eight operators counted heterotrophic plate counts from 5-30 plates (**Supplemental Figure S1**). Both the dispersion of plate counts between operators and repeat counts performed by the same operator were evaluated. While the intralaboratory uncertainty of counting was determined to be 15%, the personal uncertainty of counting varied from 2% to 11% (**Table 7**).

**Table 7.**
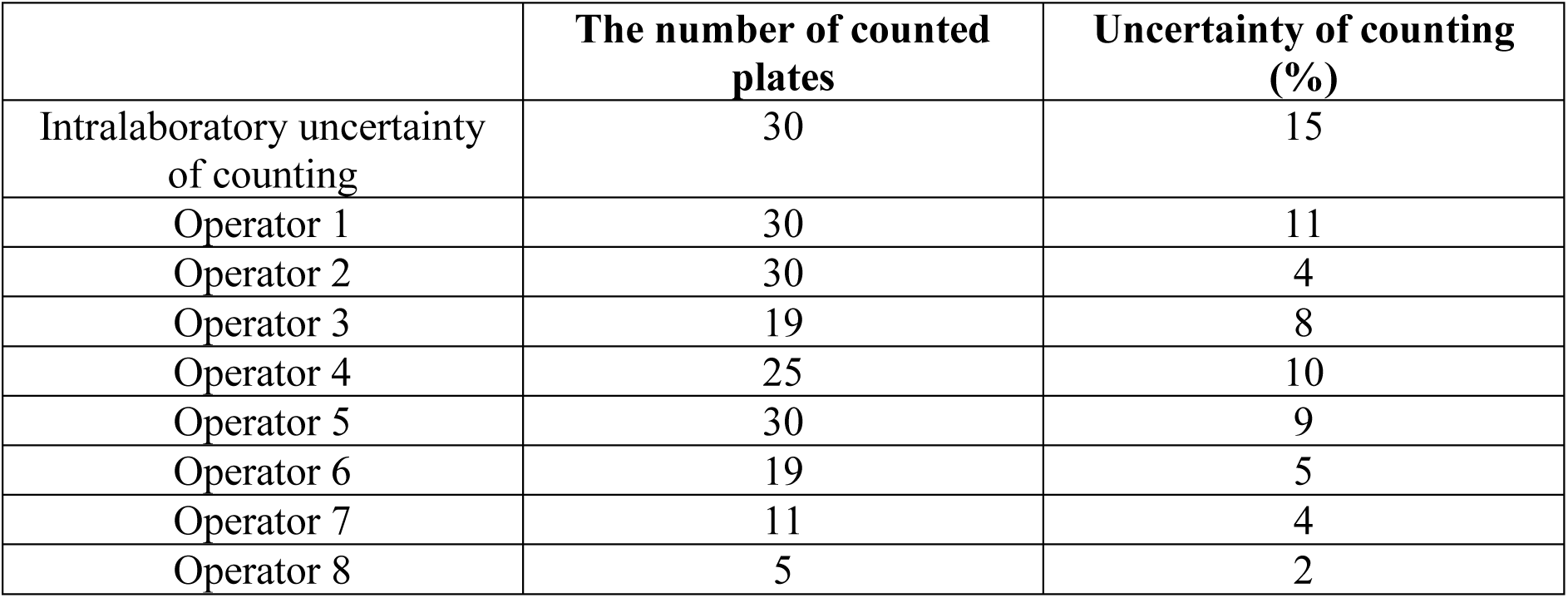
Uncertainty of counting in HPC determinations with R2A method.

#### 3.4.6. Robustness of incubation time and temperature

To determine the robustness of incubation time, two operators calculated heterotrophic plate counts from 39 plates after incubation of 160 hours, 164 hours and 168 hours. First, the relative difference of the heterotrophic plate counts from incubation of 164 hours were compared to heterotrophic plate counts from incubation of 160 hours and 168 hours according to ISO 17994:2014. The relative recovery of HPC did not differ between incubation of 164 hours and 160 hours or 164 hours and 168 hours. After that, HPC from incubation of 168 hours were compared to HPC from incubation of 160 hours. The relative recovery between incubation of 160 hours and 168 hours was not different as for both operators, the lower limit (XL) of the relative difference was more than 0% and the upper limit (XU) of the relative difference was less than +10% (**Table 8a**; Wilcoxon: p < 0.05). To determine the robustness of incubation temperature, two operators calculated heterotrophic plate counts from 21 plates after incubation at 20.5 °C and 23.5 °C. There was an average 17% difference with a 95% confidence interval [-5.5; −29] in relative recovery of HPC from incubation at 20.5 °C compared to incubation at 23.5 °C according to ISO 17994:2014 (**Table 8b**).

**Table 8.** Mean, standard deviation, lower limit (XL) and upper limit (XU) of relative difference in percentage (%) between a) incubation of 160 hours and 168 hours and b) incubation at 20.5 °C and 23.5 °C.

|  | Operator 1 | Operator 2 | Both operators |
| --- | --- | --- | --- |
| *a) Relative difference between 160 h and 168 h (N=39)* |  |  |  |
| Mean | 5.1 | 2.3 | 3.7 |
| Standard deviation | 8.4 | 6.7 | 7.7 |
| The number of plates | 39 | 39 | 78 |
| Lower limit (XL) | 2.4 | 0.1 | 2.0 |
| Upper limit (XU) | 7.8 | 4.4 | 5.4 |
| *b) Relative difference between 20.5 °C and 23.5 °C (N=21)* |  |  |  |
| Mean | -14.3 | -20.2 | -17.2 |
| Standard deviation | 39.9 | 36.6 | 38.0 |
| The number of plates | 21 | 21 | 42 |
| Lower limit (XL) | -31.7 | -36.2 | -29.0 |
| Upper limit (XU) | 3.2 | -4.2 | -5.5 |

### 3.5. Acceptable storage time between sampling and initiation of the R2A analysis

A total of 198 samples were analysed to determine acceptable storage time between sampling and initiation of the R2A analysis (**Figure 3**). The p-value of 0.373 from all data corresponds to a non-significant difference between colony counts from samples analysed at 12 hours and 24 hours after sampling. The data processing using ISO 17994 standard approach indicated an inconclusive case, close to a slight tendency towards higher results for the 24 hours (relative difference of 10,1% with a 95% confidence interval [-0.1; 20.3]). For this reason, the data from each laboratory were examined (**Supplemental Table S4**). 12-hour and 24-hour results from laboratories 2 and 8 were significantly different at a 5% risk, and the results from laboratory 4 were different at a 1% risk. A significant relative difference between colony counts from samples analysed at 12-hour and 24-hour was observed. Laboratories 2 and 8 show 24-hour results that are higher than 12-hour results. On the contrary, 24-hour results from laboratory 4 are lower than 12-hour results. Additional data processing was performed by examination of the type of water, and media suppliers, but no explanation for the results obtained by laboratory 2 and 8 (tendency towards higher results for the 24 hours) and laboratory 4 (tendency towards higher results for the 12 hours) was obtained.

**Figure 3.**
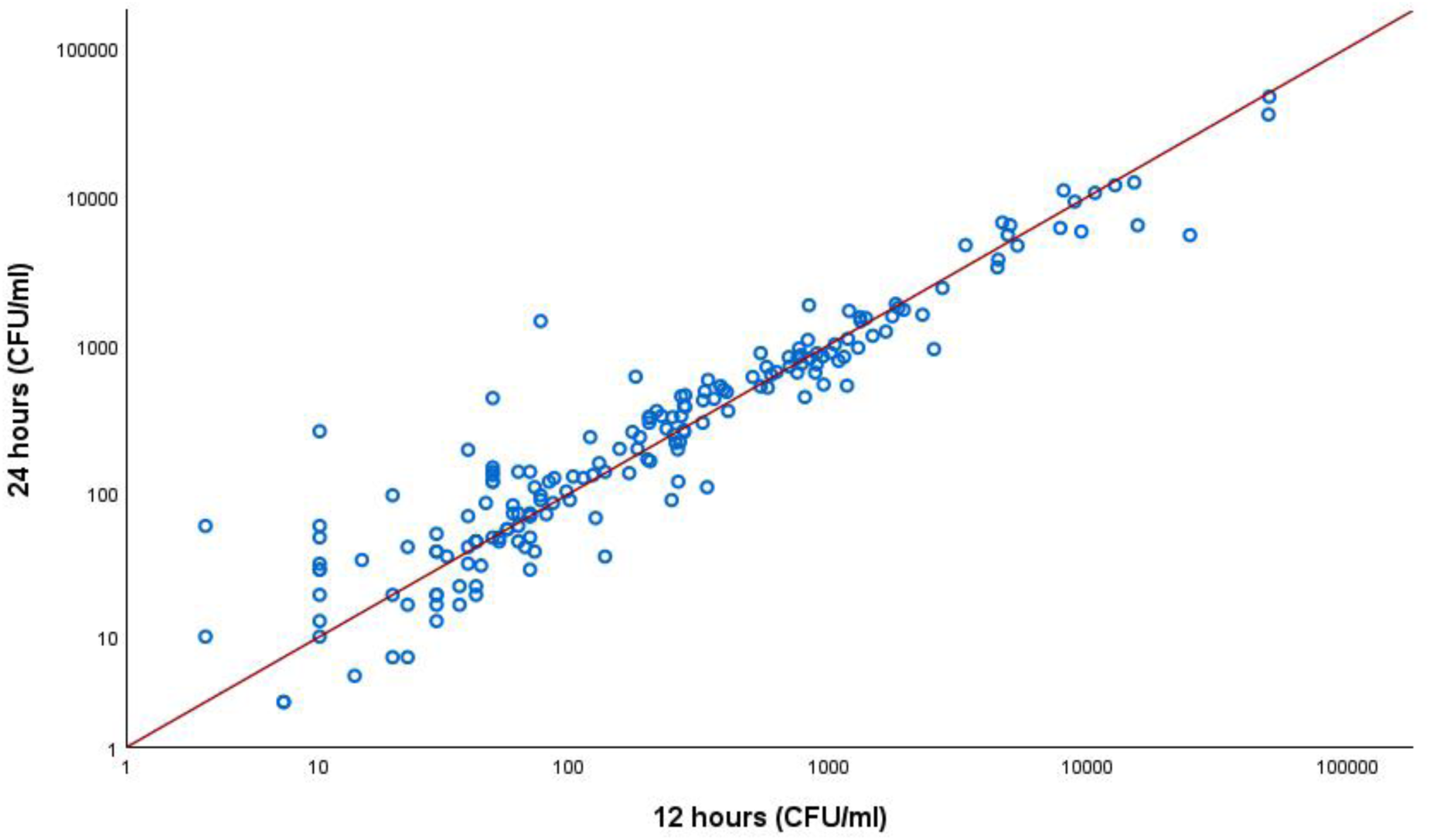
Paired colony counts of water samples on R2A medium (CFU/ml) inoculated less than 12 hours after sampling and at least 24 hours after sampling in 13 laboratories (N=198). Straight line represents the case when the colony counts are equal after 12 h and 24 h (x=y).

### 3.6. Control strains for quality assurance of the R2A medium

For all tested control strains, the productivity was consistently sufficient as it exceeded the lower limit (0.7) of the ISO 11133:2014 in each testing round. However, colonies of *Sphingomonas echinoides* (DSM 1805) were difficult to count on R2A and for this reason, this strain was omitted from further use and is not included in the standard method ISO 13647 (ISO 13647, 2026). In both laboratories, productivity of *Pseudomonas fluorescens* (WDCM 00115) and *Sphingomonas paucimobilis* (WDCM 00232) exceeded the upper limit of the productivity (1.4) of the ISO 11133:2014 during certain testing rounds (**Supplementary Table S5**).

## 4. Discussion

International standard method ISO 6222 from year 1999 is based on nutrient rich Tryptone Yeast Extract Agar medium and has been used for HPC determinations at 22 °C in 3 days or at 36 °C in 2 days. Previously, there has been national methods, but no international standard method for the use of a more sensitive Reasoner’s 2 Agar (R2A) medium that has a low-nutrient composition and long incubation time (7 days at 22 °C). This work enabled the international standardization of the R2A method (a new ISO 13647) and produced performance characteristics for R2A medium in enumeration of HPC from water by spread plate inoculation.

Determining of the performance characteristics for HPC methods differ considerably from other indicator microorganisms for water quality investigations. Namely, for example *Escherichia coli* and coliform bacteria colony counting (Lange et al., 2013) and enumeration of intestinal enterococci (Tiwari et al., 2018) is based on the use of selective culture media for which false negative rates as well as specificity, selectivity and efficiency can be determined (ISO 13843, 2017). Such determination is impossible for HPC that is based on counting all colonies from non-selective plates.

This paper describes the performance characteristics of R2A medium (Reasoner & Geldreich, 1985). While ISO 13647 (2026) specifies the use of spread plate technique, for certain purposes, especially when waters with very low nutrient content such as de-ionised, distilled or reverse osmosis waters are tested, R2A medium can be used with pour plate technique and membrane filtration techniques (ISO 15883; ISO 23500-3; ISO 23500-5). Moreover, there are other formulations available, e.g. R3A medium (Brözel & Cloete, 1992; Reasoner & Geldreich, 1985) that can be suitable for certain applications. Also, the storage conditions and time of the prepared solid agar medium plates affect the colony counts. In this study, the productivity ratio exceeded the recommended 1.4 (ISO 11133, 2014) at same testing times, tentatively due to the fact that the reference medium was prepared much earlier than the test medium.

R2A method specified herein and to be published as a new international standard method ISO 13647 (2026) provides a culture-dependent method that is a convenient tool to be further used by water companies as a relatively low-cost way of monitoring biostability and detecting general microbial failures in the system. It can be used for the assessment of water quality before, during and after the drinking water treatment processes and monitoring of integrity of the drinking water distribution systems. However, HPC determination are only representative of a limited and specific fraction of microbial communities in water samples (Reynolds & Fricker, 1999). For other purposes, there is nowadays a wide array of molecular techniques to determine the composition and functions of the drinking water and biofilm communities (Douterelo et al., 2014; Gomez-Alvarez et al., 2023), and other methods to evaluate total biomass on water, such as flowcytometry (Hammes et al., 2008).

The determined of uncertainty of colony counting in the study described herein was higher than the recommended maximum of 10% (ISO 13843, 2017), a feature affecting greatly on the performance of the R2A spread plating method and that has to be taken into account when interpreting the results. For example, the high uncertainty of the counting affect the robustness and resulted in a decision to accept a relatively wide tolerance for the range of accepted incubation temperatures, being 22 ± 2 °C in ISO 13647 (2026), although it is evident that incubation temperature does have an effect on the colony counts. The follow-up applications of ISO 13647 should also note that in this study the sample size for determinations of upper limit of counting, relative recovery, repeatability and reproducibility was lower than recommended in ISO 13843 (2017). Furthermore, the samples analysed in the single laboratory validation described herein originated from a relatively small geographical area and each laboratory must determine the performance characteristics at their premises with the local samples in their verification study prior the use of ISO 13647 (2026).

Our study verified the early observations that spread plating result in higher plate counts that pour plating technique (reviewed by Reasoner, 2004), however with the cost that also the dispersion between parallel plates is relatively high when spread plate technique is employed. The explanation to the higher colony counts after spread plating compared to plate counting could rely on the fact that the spread plate technique causes no heat shock to the bacterial cells in the sample compared to the mixing warm agar medium on a petri dish with the sample in the pour plate technique. Further, when the spread plate technique is used, all colonies are on top of the agar surface where they can be distinguished readily from particles and bubbles.

## Author contribution statement

**Kristiina Valkama:** Data curation, Formal analysis, Investigation, Visualization, Writing - original draft. **Anna Pursiainen:** Investigation, Methodology, Supervision, Writing - review & editing. **Olivier Molinier:** Formal analysis, Methodology, Supervision, Validation, Writing - review & editing. **Sunna Nikodemus:** Investigation, Visualization, Writing - review & editing. **Eric Pierlot:** Investigation, Methodology, Resources, Validation, Visualization, Writing - review & editing. **Tarja Pitkänen:** Conceptualization, Funding acquisition, Methodology, Project administration, Supervision, Writing - original draft.

## Acknowledgements

The authors gratefully acknowledge Tarja Rahkonen from the Finnish Institute for Health and Welfare (THL) for her technical assistance, together with water microbiology experts and the personnel of water laboratories worldwide who participated on this study and contributed to the data production. This work was conducted in collaboration with members of working groups 27 and 25 in the International Organization for Standardization (ISO) Technical Committee ISO/TC 147, Water quality, Subcommittee SC 4, Microbiological methods, who all are warmly acknowledged. The work was supported by the Finnish Standards Association (SFS) and was conducted in collaboration with University of Eastern Finland and with VEMISTA, a national mirror committee for standardization of water microbiology in Finland.

## Notes

### Competing Interest Statement

The authors have declared no competing interest.

