## Supplemental material for "Performance of Reasoner’s 2 Agar (R2A) medium in enumeration of culturable microorganisms in water intended for human consumption"

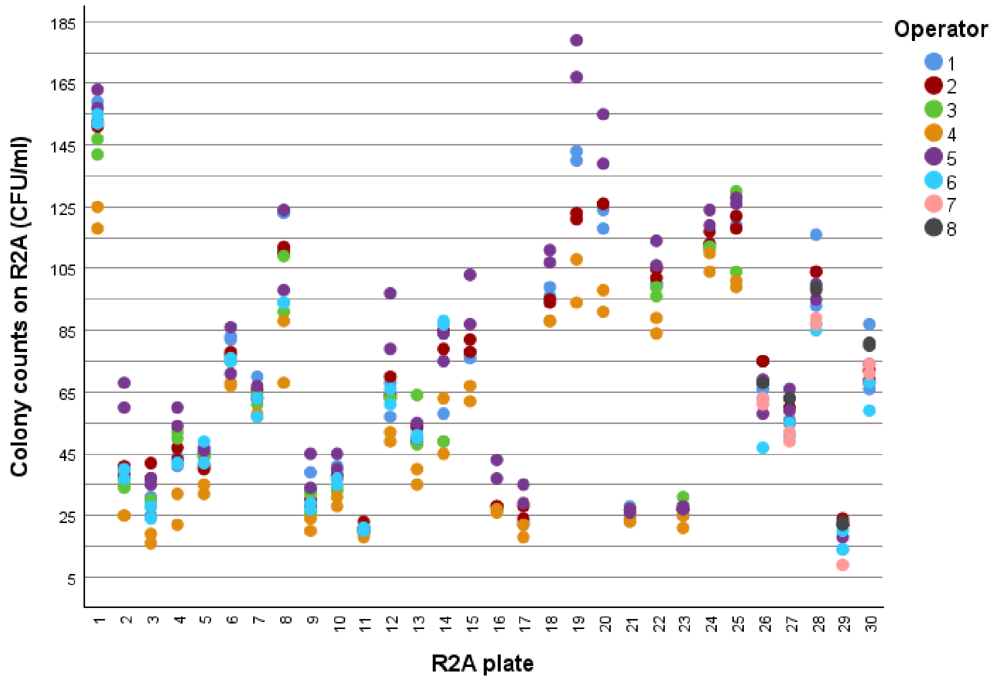


**Supplemental Figure S1.** Colony counts on 30 R2A plates (CFU/ml). The HPC determined by operators 1-8 are shown in different colours.

**Supplemental Table S1.** Samples analysed by the 11 laboratories, their arithmetical means, standard deviations and arithmetic mean of operational variance ($u_o^2$) using spread plate and pour plate techniques.

| **Laboratory** | **Number of samples** | **Spread plate technique** | | **Pour plate technique** | |
|---|---|---|---|---|---|
| | | **Arithmetic mean ± standard deviation** | **Arithmetic mean of relative operational variance** | **Arithmetic mean ± standard deviation** | **Arithmetic mean of relative operational variance** |
| 1 | 15 | 32 ± 39 | 0.005 | 30 ± 38 | 0.026 |
| 2 | 23 | 88 ± 56 | 0.116 | 30 ± 29 | -0.006 |
| 3 | 33 | 72 ± 38 | 0.008 | 34 ± 29 | -0.005 |
| 4 | 18 | 71 ± 53 | 0.023 | 46 ± 39 | -0.007 |
| 5 | 16 | 66 ± 46 | -0.005 | 33 ± 34 | 0.028 |
| 6 | 3 | 147 ± 20 | 0.017 | 160 ± 20 | -0.003 |
| 7 | 13 | 62 ± 51 | 0.026 | 53 ± 45 | -0.020 |
| 8 | 5 | 41 ± 34 | -0.005 | 16 ± 15 | -0.036 |
| 9 | 22 | 61 ± 40 | -0.002 | 43 ± 29 | -0.003 |
| 10 | 22 | 59 ± 40 | -0.006 | 37 ± 33 | -0.011 |
| 11 | 20 | 47 ± 33 | 0.092 | 47 ± 37 | 0.041 |
| **Total** | **190** | **65 ± 47** | **0.028** | **40 ± 37** | **0.002** |

**Supplemental Table S2.** Heterotrophic plate counts from water samples studied by using different diluents at inoculation time points 0 min and 45 min after preparation of the 10-fold dilution series. Duplicate spread plating of samples at each dilution level was performed with each diluent at both time points.

| | **Sterile water** | | **Physiological saline** | | **Phosphate buffer** | | **Peptone water** | | **Peptone saline solution** | |
|---|---|---|---|---|---|---|---|---|---|---|
| **Time point (min)** | 0 | 45 | 0 | 45 | 0 | 45 | 0 | 45 | 0 | 45 |
| **Sum of colonies** | 916 | 828 | 854 | 669 | 859 | 761 | 836 | 913 | 776 | 644 |
| **Arithmetic mean** | 51 | 46 | 47 | 37 | 48 | 42 | 26 | 51 | 43 | 36 |
| **Median** | 19 | 20 | 19 | 15 | 21 | 20 | 20 | 27 | 25 | 15 |
| **Standard deviation** | 58 | 50 | 55 | 43 | 52 | 45 | 51 | 51 | 43 | 47 |

**Supplemental Table S3.** Sum of colonies observed on the R2A plates and descriptive statistics of the plate counts when comparing a light source above and below using 2.25x or 3x magnification and a stereo microscope (N=24 plates).

| | **2.25x, light above** | **2.25x, light below** | **3x, light above** | **3x, light below** | **Stereo microscope** |
|---|---|---|---|---|---|
| **Sum of colonies** | 3798 | 3870 | 3212 | 3692 | 3790 |
| **Arithmetic mean** | 46 | 47 | 39 | 45 | 47 |
| **Median** | 30 | 27 | 24 | 26 | 28 |
| **Standard deviation** | 39 | 42 | 37 | 40 | 41 |

**Supplemental Table S4.** Statistical analysis of all data and of each laboratory from samples analysed at 12 hours and 24 hours after sampling.

| | | Wilcoxon test | | ISO 17994 (adapted) | | |
|---|---|---|---|---|---|---|
| **Laboratory** | **Number of exploitable paired data** | **p-value** | **Conclusion** | **Relative difference** | **95% confidence interval** | **Conclusion** |
| 1 | 24 | 0.903 | non-significant | 2.9% | [-11.8; 17.6] | inconclusive[c] /not different[d] |
| 2 | 10 | 0.021 | *5% | 27.6% | [13.7; 41.4] | tendency towards higher results for the 24 h |
| 3 | 12 | 0.929 | non-significant | -1.4% | [-14.5; 11.7] | inconclusive[c] /not different[d] |
| 4 | 18 | 0.003 | **1% | -25.9% | [-41.9; -9.9] | tendency towards higher results for the 12 h |
| 5 | 28 | 0.400 | non-significant[a] | -1.4% | [-27.1; 24.3] | inconclusive |
| 6 | 23 | 0.175 | non-significant[a] | 48.7% | [4.9; 92.4] | tendency towards higher results for the 24 h |
| 7 | 7 | 0.236 | non-significant | not estimated | not estimated | not estimated |
| 8 | 10 | 0.013 | *5% | 31.9% | [10.5; 53.3] | tendency towards higher results for the 24 h |
| 9 | 3 | not estimated | not estimated | not estimated | not estimated | not estimated |
| 10 | 5 | 0.500 | non-significant[a] | not estimated | not estimated | not estimated |
| 11 | 19 | 0.255 | non-significant[a] | 28.4% | [-9.0; 65.7] | inconclusive |
| 12 | 1 | not estimated | not estimated[a] | not estimated | not estimated | not estimated |
| 13 | 8 | 0.735 | non-significant[a] | not estimated | not estimated | not estimated |
| **Total number of laboratories** | **Total number of exploitable paired data** | **All data Wilcoxon test p-value** | | **All data ISO 17994** | | |
| | | **p-value** | **Conclusion** | **Relative difference** | **95% confidence interval** | **Conclusion** |
| 13 | 168 | 0.373 | non-significant | 10.1% | [-0.1; 20.3] | inconclusive case, close to a slight tendency towards higher results for the 24 h |

*Significant test at 5% risk of error. **Significant test at 1% risk of error. [a]Number of sample pairs falling outside the agreed range of storage time prior analysis: laboratory 5 (1 sample); laboratory 6 (7 samples); laboratory 7 (1 sample); laboratory 12 (7 samples). [b]The trend is based on the presence of two extreme points, in the absence of which it is no longer significant. [c]For drinking water (± 10%). [d]For environmental waters (± 20%).

**Supplementary Table S5.** Productivity of *Pseudomonas fluorescens* (WDCM 00115), *Bacillus subtilis subsp. spizizenii* (WDCM 00003), *Sphingomonas paucimobilis* (WDCM 00232) and *Sphingomonas echinoides* (DSM 1805). NA = not analysed.

| | Productivity (Pr) | | | | | | | |
|---|---|---|---|---|---|---|---|---|
| | Laboratory 1 | | | | Laboratory 2 | | | |
| **Control strain** | **Range** | **Arithmetic mean** | **Standard deviation** | **N** | **Range** | **Arithmetic mean** | **Standard deviation** | **N** |
| *Pseudomonas fluorescens* (WDCM 00115) | 0.71-1.75 | 1.09 | 0.23 | 20 | 1.13-1.58 | 1.37 | 0.19 | 3 |
| *Bacillus subtilis subsp. spizizenii* (WDCM 00003) | 0.77-1.25 | 0.99 | 0.15 | 27 | 1.04-1.09 | 1.06 | 0.02 | 3 |
| *Sphingomonas paucimobilis* (WDCM 00232) | 0.90-1.65 | 1.19 | 0.23 | 19 | 1.09-2.32 | 1.84 | 0.54 | 3 |
| *Sphingomonas echinoides* (DSM 1805) | 0.90-1.05 | 1.00 | 0.07 | 4 | NA | NA | NA | NA |
